# A Polygenic Score for Higher Educational Attainment is Associated with Larger Brains

**DOI:** 10.1101/287490

**Authors:** Maxwell L. Elliott, Daniel W Belsky, Kevin Anderson, David L. Corcoran, Tian Ge, Annchen Knodt, Joseph A. Prinz, Karen Sugden, Benjamin Williams, David Ireland, Richie Poulton, Avshalom Caspi, Avram Holmes, Terrie Moffitt, Ahmad R Hariri

**Affiliations:** Department of Psychology & Neuroscience, Duke University Box 104410, Durham, NC 27708, USA; Department of Population Health Sciences, Duke University School of Medicine, Box 3003, Durham, NC 27710, USA; Social Science Research Institute, Duke University, Box 90989, Durham, NC 27708, USA; Department of Psychology, Yale University, New Haven, CT 06511, USA; Center for Genomic and Computational Biology, Duke University Box 90338, Durham, NC 27708, USA; Athinoula A. Martinos Center for Biomedical Imaging, Massachusetts General Hospital, Harvard Medical School, Charlestown, MA 02129, USA; Psychiatric and Neurodevelopmental Genetics Unit, Center for Genomic Medicine, Massachusetts General Hospital, Boston, MA 02114, USA; Stanley Center for Psychiatric Research, Broad Institute of MIT and Harvard, Cambridge, MA, USA; Dunedin Multidisciplinary Health and Development Research Unit, Department of Psychology, University of Otago, 163 Union St E, Dunedin, 9016, NZ; Social, Genetic, & Developmental Psychiatry Research Centre, Institute of Psychiatry, Psychology, & Neuroscience, King’s College London, De Crespigny Park, Denmark Hill, London SE5 8AF, UK; Department of Psychiatry & Behavioral Sciences, Duke University School of Medicine, Durham, NC 27708, USA

## Abstract

People who score higher on intelligence tests tend to have larger brains. Twin studies suggest the same genetic factors influence both brain size and intelligence. This has led to the hypothesis that genetics influence intelligence partly by contributing to development of larger brains. We tested this hypothesis with molecular genetic data using discoveries from a genome-wide association study (GWAS) of educational attainment, a correlate of intelligence. We analyzed genetic, brain imaging, and cognitive test data from the UK Biobank, the Dunedin Study, the Brain Genomics Superstruct Project (GSP), and the Duke Neurogenetics Study (DNS) (combined N=8,271). We measured genetics using polygenic scores based on published GWAS. We conducted meta-analysis to test associations among participants’ genetics, total brain volume (i.e., brain size), and cognitive test performance. Consistent with previous findings, participants with higher polygenic scores achieved higher scores on cognitive tests, as did participants with larger brains. Participants with higher polygenic scores also had larger brains. We found some evidence that brain size partly mediated associations between participants’ education polygenic scores and their cognitive test performance. Effect-sizes were larger in the population-based UK Biobank and Dunedin samples than in the GSP and DNS samples. Sensitivity analysis suggested this effect-size difference partly reflected restricted range of cognitive performance in the GSP and DNS samples. Recruitment and retention of population-representative samples should be a priority for neuroscience research. Findings suggest promise for studies integrating GWAS discoveries with brain imaging data to understand neurobiology linking genetics with individual differences in cognitive performance.

## Introduction

People who score higher on tests of intelligence tend to have larger brains, as measured by ex-vivo brain weight and in-vivo magnetic resonance imaging (MRI)^1–4^. Twin studies indicate this relationship partly reflects genetic factors that influence both brain size (i.e., volume) and intelligence^5–8^. These findings suggest the hypothesis that one path through which genetic differences between people influence individual differences in intelligence is by contributing to the development of larger brains. This hypothesis can now be tested using molecular genetic data.

A recent genome-wide association study (GWAS) of educational attainment identified dozens of genetic variants that showed substantial enrichment for genes expressed during brain development^9^. Follow-up studies further identified associations between an aggregate measure of GWAS-discovered influences on education, called a polygenic score, and intelligence, including in young children who had not yet entered school^10,11^. These findings implicate brain development and intelligence in the pathway connecting people’s genetics to their educational outcomes. Further, GWAS research has discovered polygenic variants associated with brain size (inferred through intracranial volume)^12^ that also overlap with variants associated educational attainment^9^. Now, studies are needed to test if genetics discovered in GWAS of education are associated with in-vivo intermediate phenotypes, like brain size, that could constitute a biological pathway linking genetic variation to differences in intelligence and educational attainment.

We analyzed data from four imaging genetics studies from the United Kingdom (UK Biobank), New Zealand (Dunedin Study), and the United States (Brain Genomics Superstruct Project (GSP) and Duke Neurogenetics Study), including 8,271 participants, to test associations among a polygenic score for educational attainment, cognitive test performance, and brain size. We hypothesized that, consistent with previous findings, (1) participants with higher education polygenic scores would have higher cognitive test scores; and (2) that participants with larger brains as measured by total brain volume would have higher cognitive test scores. We further posed the novel hypotheses that (3) participants with higher education polygenic scores would have larger brains and that brain size would mediate the association between the education polygenic score and cognitive test performance. We combined results across our four imaging genetics datasets using random-effects meta-analysis. We also examined heterogeneity between the datasets under the hypothesis that effect-sizes might differ between the population-based UK Biobank and Dunedin Study samples and the GSP and DNS samples, for which range in cognitive performance is more restricted.

## Methods

### Participants

We analyzed data from European-descent participants in the United Kingdom-based UK Biobank^13,14^ a population-based volunteer sample (N=6117), the New Zealand-based Dunedin Study, a birth cohort (N=476)^15^, and two studies in the United States consisting primarily of university students, the Brain Genomics Superstruct Project^16^ (GSP, N=1163), and the Duke Neurogenetics Study^17^ (DNS, N=515). Sample sizes reflect participants with available structural MRI, cognitive testing, and genetic data (Table 1). Samples are described in detail in the supplement.

**Table 1.** Samples and measures included in analysis. Polygenic scores for all samples were computed based on the most recent GWAS of educational attainment^9^ following established methods.

| Sample | Cognitive Test | Total Brain Volume (cm<sup>3</sup>) |
| --- | --- | --- |
| *United Kingdom Biobank (UK Biobank)*<sup>13</sup>: An ongoing general population-based cohort of volunteers that was recruited from the UK National Health Service records beginning in 2006.<br>N = 6117<br>53% female.<br>Age M = 61.39, SD = 7.00 | 13 verbal-numeric reasoning puzzles completed during a 2-minute time test<sup>21</sup>.<br>Scored as number of correct responses.<br>M = 6.95, SD = 2.10 | Total brain volume was derived from T1 weighted structural MRI images processed with Sienax<sup>25</sup><br>M = 1170.86, SD = 111.14 |
| *Dunedin Multidisciplinary Health and Development Study (DMHDS)*<sup>15</sup>: a population representative birth cohort born 1972-3 in Dunedin, New Zealand. Note: Here we report the available N, as of 2.2018, while data collection is ongoing.<br>N = 476<br>51% female<br>Intelligence testing age = 38, MRI testing age = 45 | Wechsler Adult Intelligence Scale-IV (WAIS-IV):<sup>22</sup><br>Scored against a population norm with mean of 100 and standard deviation of 15.<br>M = 99, SD = 15 | Total brain volume was derived from the recon-all pipeline in Freesurfer<sup>26</sup> using T1 and T2 weighted structural MRI images.<br>M = 1218.32, SD = 121.36 |
| *Brain Genomics Superstruct Project (GSP)*<sup>16</sup>: a convenience sample of Boston area healthy volunteers primarily recruited from local universities and medical centers.<br>N = 1163<br>53% Female<br>Age M = 22.23, SD = 5.53 | Shipley Institute of Living Scale<sup>23</sup>.<br>Scored against a population norm with mean of 100 and standard deviation of 15.<br>M = 113, SD = 9 | Total brain volume was derived from the recon-all pipeline in Freesurfer<sup>26</sup> using T1 and T2 weighted structural MRI images.<br>M = 1174.58, SD = 110.64 |
| *Duke Neurogenetics Study (DNS)*: A convenience sample of university students primarily from Duke University.<br>N = 515<br>53% female<br>Age M = 20.26, SD = 1.20 | Matrix reasoning and vocabulary subtests of the Wechsler Abbreviated Scale of Intelligence<sup>24</sup> (WASI)<br>Scored against a population norm with mean of 100 and standard deviation of 15.<br>M = 124, SD = 7 | Total brain volume was derived from the recon-all pipeline in Freesurfer<sup>26</sup> using T1 weighted structural MRI images.<br>M = 1162.40, SD = 110.34 |

### Education Polygenic Score

We computed our polygenic score based on GWAS of educational attainment rather than GWAS of cognitive performance because educational attainment is a proxy phenotype for cognitive performance^18^ and the polygenic score for educational attainment is more predictive of cognitive performance than polygenic scores from GWAS of cognitive performance^19^. Education polygenic scores were computed from genome-wide single-nucleotide polymorphism (SNP) data based on GWAS results published by the Social Science Genetics Association Consortium^9^ following methods described by Dudbridge^20^ according to the procedure used in our previous work^10^. Briefly, for each study, we matched SNPs in the study’s genetic database with published educational attainment GWAS results^9^. We then multiplied the education-associated allele of each SNP by the GWAS-estimated effect-size and computed the average of these products across all SNPs. Polygenic scores were standardized within each study to have M=0, SD=1 for analysis.

### Cognitive Performance

Cognitive performance was measured in the UK Biobank using 13 reason and logic puzzles^21^ and in the Dunedin Study, GSP, and DNS studies using intelligence tests (the Wechsler Adult Intelligence Scale (WAIS-IV)^22^ in the Dunedin Study, the Shipley Institute of Living Scale^23^ in GSP and the Wechsler Abbreviated Scale of Intelligence (WASI)^24^ in the DNS).

### Total Brain Volume

Total brain volume was measured from high resolution, T1-weighted MRI images. In the UK Biobank total brain volume was estimated using SIENAX^25^. In the Dunedin Study, GSP, and DNS studies, images were processed using the Freesurfer processing pipeline^26^.

### Statistical Analyses

We tested associations using linear regression models. Models were adjusted for sex. Models including the polygenic score were adjusted for the first 10 principal components estimated from the genome-wide SNP data to account for any residual population stratification within the European-descent samples analyzed^27^. Models of UK biobank and GSP data were adjusted for age. (The Dunedin Study is a single-year birth cohort and DNS participants vary in age by only by 1-2 years.). In addition to age, models in the GSP were also adjusted for scanner, console version and head coil (12 versus 32 channel) because the GSP was collected across multiple sites. Analyses of individual studies were conducted in R (version 3.4.0). Linear regressions were performed using the lm function. Mediation analyses were performed using a system of equations approach^28^ implemented with the *mediation* package^29^ in R, using nonparametric bootstrapping with 1000 iterations. We combined estimates across studies using random effects meta-analysis^30^ implemented using STATA (version 15).

## Results

### Participants with higher polygenic scores performed better on cognitive tests

As anticipated, participants with higher polygenic scores performed better on cognitive tests. Meta-analysis estimated the cross-study effect size as r=.18 (p<.001; 95% CI [.12, .24]) with evidence of heterogeneity in effect sizes across studies (I-squared 80%, p=.002). Effect sizes were statistically significant in UK Biobank (r=.20, p<.001), Dunedin Study (r=.28, p<.001) and GSP (r=.19, p<.001) but not in the DNS (r=.05, p=.220).

### Participants with larger brains had higher cognitive test scores

We next tested if participants with larger brains performed better on cognitive tests. As anticipated, participants with larger brains (i.e., those with higher total brain volume) performed better on cognitive tests. Meta-analysis estimated the cross-study effect size as r=.20 (p<.001; 95% CI [.12, .28]) with evidence of heterogeneity in effect sizes across studies (I-squared=75.8%, p=.006). Effect-sizes were statistically significant in all studies (UK Biobank r=.21, p<.001; Dunedin Study r=.35, p<.001; GSP r=.12, p=.002; DNS r=.16, p=.004).

### Participants with higher polygenic scores for educational attainment had larger brains in two samples

Finally, we tested if participants with higher polygenic scores tended to have larger brains. Meta-analysis estimated the cross-study effect-size as r=.05 (p=.002; 95% CI [.02, .09]). The test for evidence of heterogeneity in effect sizes across studies was not statistically significant at the alpha=.05 level (I-squared=51.9%, p=.101). Participants with higher polygenic scores had larger brains in the UK Biobank (r=.08, p<.001) and the Dunedin Study (r=.08, p=.033). Effect-sizes were smaller and not statistically significant in the GSP r=.02, p=.380 and DNS r=.04, p=.288.

### Brain size was a weak mediator of the polygenic-score associations with cognitive test scores in two study samples

To test the hypothesis that larger brains mediated the polygenic score association with intelligence, we used the system of equations described by Baron and Kenny^31^ and the methods described by Preacher et al.^28^ Meta-analysis estimated the cross-study indirect effect to be b=.01, 95% CI [.00, .02], p=.055, with evidence of heterogeneity in effect sizes across studies (I-squared=79.5%, p=.002). The mediation effect was statistically significant in the UK Biobank (b=.01, 95% CI [.01, .02], p < .001) and the Dunedin Study (b=.02, 95% CI [.00, .05], p=.042). We did not find evidence of a mediation effect in the GSP b=.00, 95% CI [.00, .01], p=.36) or DNS b=.01, 95% CI [-.00, .02], p=.24.

### Sensitivity analysis: Associations among polygenic scores, brain size, and cognitive test scores were attenuated in a sample of UK Biobank participants restricted to those with cognitive test scores 1 SD above the sample mean

UK Biobank and Dunedin Study participants’ polygenic scores, brain size, and cognitive test performance were positively correlated, with similar effect-sizes (Dunedin-study effect-sizes for analyses including IQ were somewhat larger, possibly reflecting greater measurement precision of the WAIS as compared to the UK Biobank reason-and-logic-puzzle test). By comparison, effect-sizes for these associations were smaller among GSP and DNS participants. To test if this difference could reflect the relatively restricted range of cognitive test performance in the GSP and DNS samples relative to the population-based UK Biobank and Dunedin samples, we conducted sensitivity analysis.

Cognitive test scores were on average, 1-1.5 SDs higher in the GSP and DNS samples as compared to the general population and 30-50% less variable, indicating restricted range (Table 1). Sensitivity analysis restricted the UK Biobank sample to participants with cognitive test scores 1 SD above the full-sample mean (i.e. scores of 9-13; n=1,401), for which the variance was approximately 45% of the full-sample variance. In the restricted-sample sensitivity analysis, associations among participants’ polygenic scores, brain size, and cognitive test performance were attenuated by roughly 1/3 to 1/2 relative to the full-sample estimates (**Supplemental Table S3**).

## Discussion

We analyzed data from four imaging genetics studies in the UK, NZ, and US to test if genetic associations with cognitive performance were mediated by differences in brain size. As anticipated, we found that participants with higher educational-attainment polygenic scores tended to score higher on tests of cognitive performance, as did those with larger brains. We also found new information, that participants with higher education polygenic scores tended to have larger brains. In mediation analysis, brain size accounted for only a small fraction of the association between participants’ educational attainment polygenic scores and their cognitive performance, and this mediation effect was statistically significant in the population-based UK Biobank and Dunedin samples, but not in the GSP and DNS samples.

Effect-size variation across the samples we analyzed followed a consistent pattern; effect-sizes were larger in the population-based UK Biobank and Dunedin Study samples than in the GSP and DNS samples. One reason for these differences may be the more restricted range of variation in cognitive performance in the GSP and DNS samples arising from, e.g. overrepresentation of university educated individuals. Such range restriction biases association estimates^32,33^ and has previously been shown to bias brain imaging research^34,35^. In the these relatively high IQ and restricted range samples, average cognitive performance was 1-1.5 standard deviations above the general-population mean and the variance was reduced by 30-50%. We conducted sensitivity analysis in a UK Biobank subsample selected to have high cognitive performance similar to the GSP and DNS samples. In this sample with restricted range of cognitive test performance, effect-sizes were attenuated by roughly 30-50%. Selective observation of high-cognitive-performance individuals in the GSP and DNS samples may have contributed to the lower effect-size estimates in these samples and to overall heterogeneity across samples in our meta-analysis.

We acknowledge limitations of our current analyses, which can be addressed in future research. First, analyses were restricted to European-descent participants. We focused on European-descent participants to match the population studied in the GWAS of educational attainment^8^. Application of GWAS results from European-descent samples to compute polygenic scores for samples of different ancestry has uncertain validity^36^. As GWAS of education and related phenotypes in non-European samples become available, replication in additional populations will be needed. Second, polygenic scores were measured with substantial error. Genetic effect-sizes thus represent lower-bound estimates. As larger-sample GWAS become available, error in polygenic score measurement will decline and effect-sizes can be expected to increase^37^. Third, total brain volume is only one route through which the genetics linked with educational attainment could affect cognitive performance. We studied this specific phenotype because it is the best-replicated neural correlate of cognitive function^2^. As more refined neural phenotypes of cognitive function are developed, including measures of cortical thickness, surface area, gyrification, and brain function, it will be important to test their potential mediating role in linking genetics with cognitive performance. Finally, we cannot rule out age differences as a potential explanation for the difference in findings between the population-based UK Biobank and Dunedin Study samples as compared to the GSP and DNS samples. UK Biobank and Dunedin Study participants were measured in midlife, whereas GSP and DNS samples primarily included young adults. Among midlife UK Biobank participants, restricting the range of cognitive performance to be similar to the GSP and DNS samples reduced effect-sizes for associations among polygenic scores, brain size, and cognitive test performance. Population-based samples including both young and midlife individuals with DNA, MRI, and cognitive testing are needed to evaluate whether genetic associations with brain volume and cognitive performance vary with age. A final concern is potential reverse causation between brain size and cognitive function. Higher cognitive ability and related educational and socioeconomic attainments may be protective of age-related decline in brain volume. Longitudinal studies with repeated measures of brain volume and cognition are needed to establish causal direction.

Within the bounds of these limitations, our findings contribute to evidence that genetics discovered in GWAS of educational attainment influence brain development and cognitive function. Bioinformatic analysis of education GWAS results have identified enrichment of variants near genes expressed in brain development, specifically neural proliferation, neural development, and dendrite formation^9^. Epidemiologic analysis of an education-GWAS-based polygenic score found that children who carried more education-associated genetic variants scored higher on cognitive tests as early as age 5 and that polygenic-score-associated differences in cognitive test scores grew larger from middle childhood through adolescence^10^. Several studies have reported that an education-GWAS-based polygenic score is predictive of cognitive test performance in adolescents and adults^11,19,38^. Here, we show that adults with higher education-GWAS-based polygenic scores have larger brains and score higher on cognitive tests as compared to peers with lower polygenic scores. Evidence for larger brains as a statistical mediator of polygenic score associations with cognitive performance was mixed in our analysis. But findings suggest promise for future neuroscientific investigation of education-linked genetics. One design to complement formal mediation analysis is gene-environment interaction analysis to test if exposures that slow brain growth or restrict brain size, e.g., Zika virus^39^, diminish associations between genetics and cognitive performance.

Our finding that genetics associated with educational and socioeconomic attainments are also related to brain volume has implications for research on effects of poverty on the developing brain. Childhood poverty exposure is associated with smaller brain volumes^40,41^. Education polygenic scores also tend to be lower in children growing up in poorer families, a gene-environment correlation that presumably reflects effects of education-linked genetics on parents economic attainments, which children inherit along with their genotypes^10,30^. Studies that include controls for education genetics could complement intervention studies^42^ to help rule out potential confounding in associations between poverty and brain development.

A challenge facing research on how genetics affect the brain is the lack of population-representative samples with available brain imaging data. Human brain-imaging research has typically been conducted in samples similar to those in the GSP and DNS whose data we analyzed^43,44^. Our findings illustrate how studies of samples pre-selected for high levels of cognitive functioning and related characteristics impose limitations on analysis of cognition-related neurobiology. Opportunities to understand the brain afforded by 21^st^ Century measurement technologies must still reckon with 20^th^ Century discoveries about selection bias^45,46^. Efforts to recruit more representative samples that reflect the full range of cognitive functioning in the population are needed.

Individual differences in cognitive performance have a partial genetic etiology^19,47^. This genetic etiology should be evident in individual differences in brain biology. As GWAS discoveries for intelligence and related traits clarify genetic etiology, follow-up in genetically-informed brain imaging studies can shed light on the neurobiological correlates of this genetic variation. Our findings encourage enthusiasm for this research, but also highlight limitations of existing data resources. Recruiting and retaining samples that are representative of the general population must be a priority in neuroscience research.

**Figure 1.**
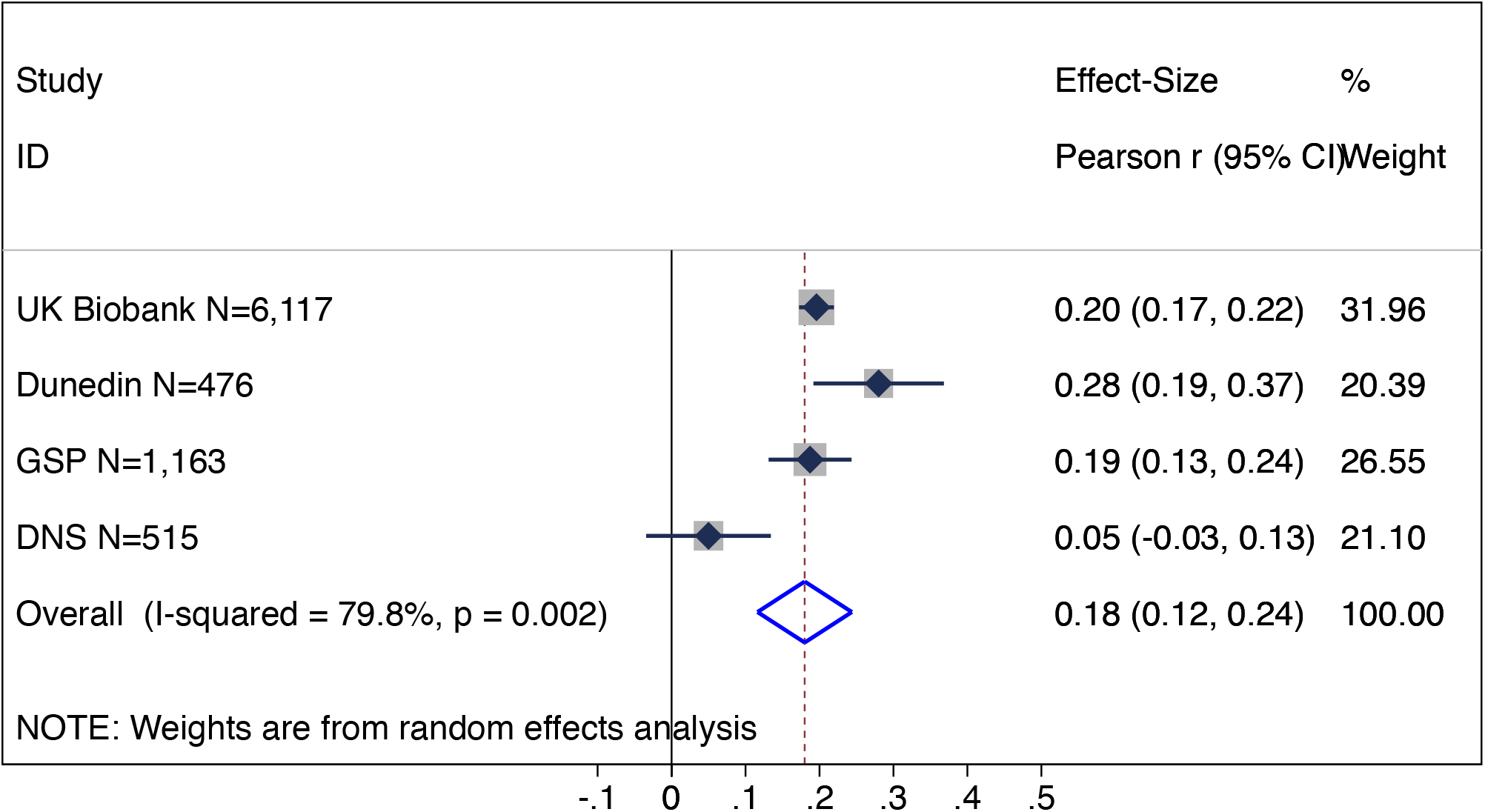
Educational attainment polygenic score associations with cognitive test scores. The figure shows a graph of effect-sizes for analyses of the UK Biobank, Dunedin Study (Dunedin), Brain Genomics Superstruct Project (GSP) and Duke Neurogenetics Study (DNS) samples (solid blue diamonds) and the cross-study effect-size estimated from random-effects meta-analysis (open blue diamond). Gray boxes around the solid-blue diamonds show the weighting of study-specific estimates in the meta-analysis (larger gray boxes indicate higher weights). 95% CIs for estimates are shown as error bars for the study-specific estimates and as the left- and right-extremes of the diamond for the meta-analysis effect-size. The meta-analysis estimate of between-study heterogeneity (I-squared) is listed to the left of the open blue diamond showing the meta-analysis effect-size. The table to the right of the effect-size graph reports values for effect-sizes, 95% CIs, and meta-analysis weights.

**Figure 2.**
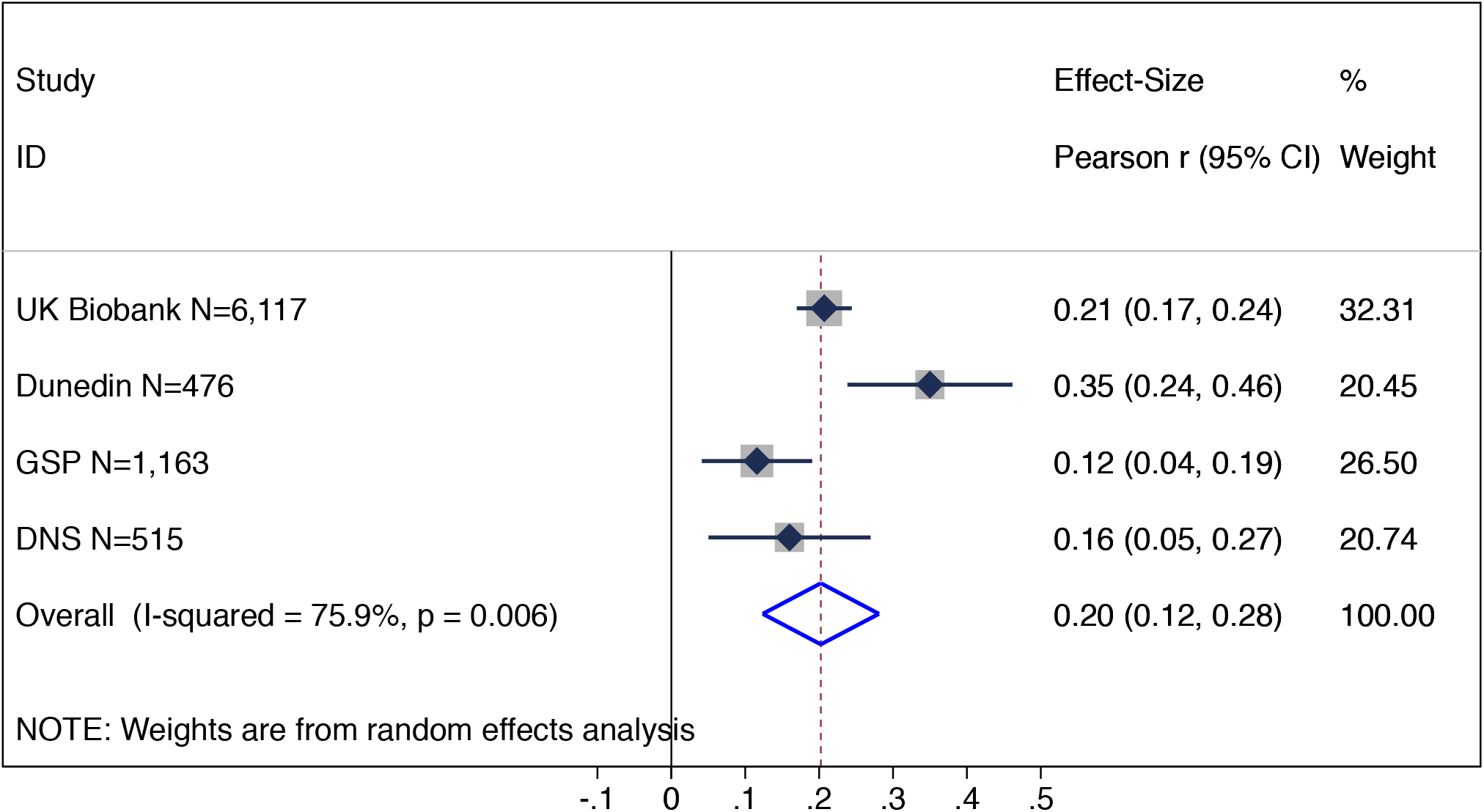
Associations between brain size and cognitive test scores. The figure shows a graph of effect-sizes for analyses of the UK Biobank, Dunedin Study (Dunedin), Brain Genomics Superstruct Project (GSP) and Duke Neurogenetics Study (DNS) samples (solid blue diamonds) and the cross-study effect-size estimated from random-effects meta-analysis (open blue diamond). Gray boxes around the solid-blue diamonds show the weighting of study-specific estimates in the meta-analysis (larger gray boxes indicate higher weights). 95% CIs for estimates are shown as error bars for the study-specific estimates and as the left- and right-extremes of the diamond for the meta-analysis effect-size. The meta-analysis estimate of between-study heterogeneity (I-squared) is listed to the left of the open blue diamond showing the meta-analysis effect-size. The table to the right of the effect-size graph reports values for effect-sizes, 95% CIs, and meta-analysis weights.

**Figure 3.**
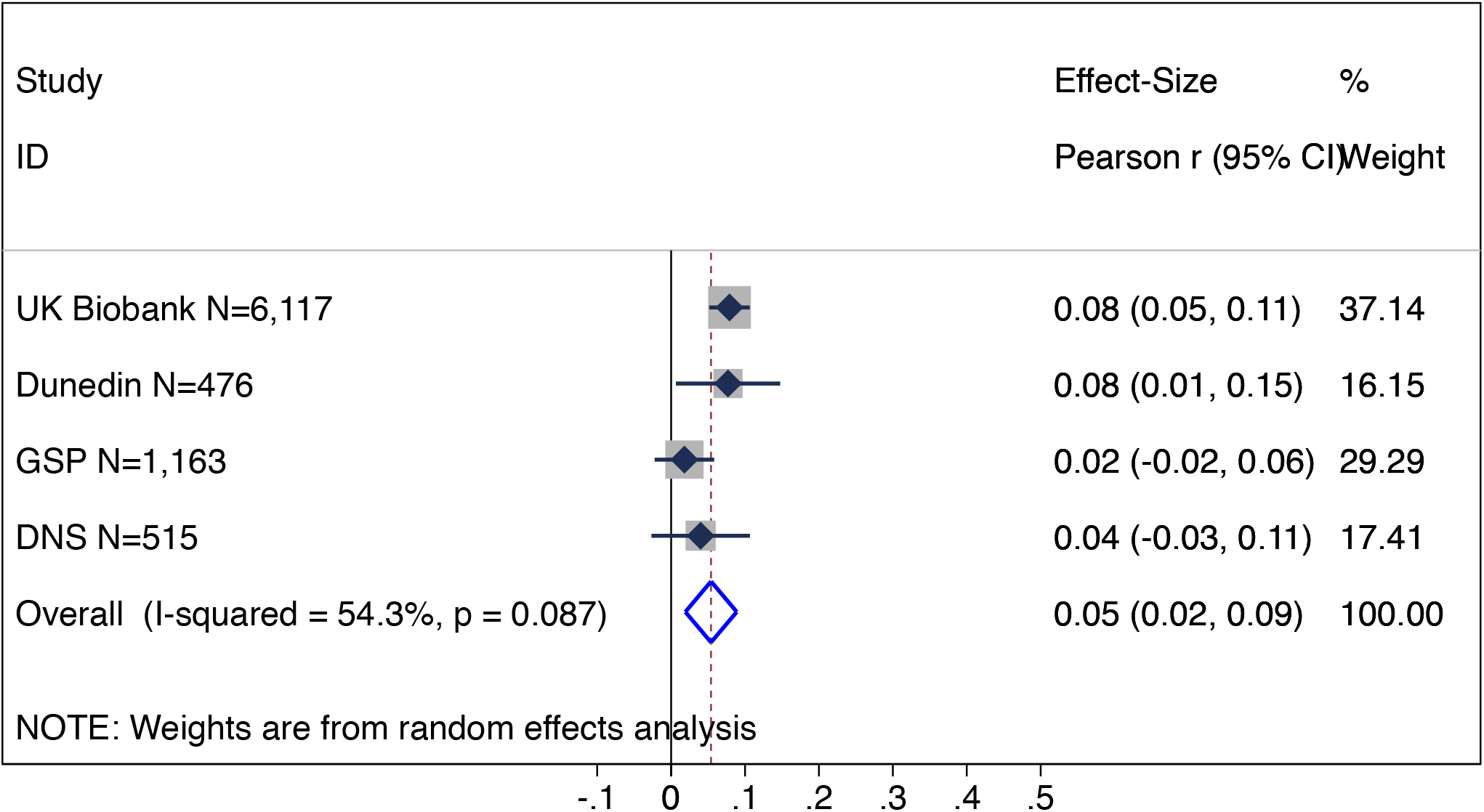
Educational attainment polygenic score associations with brain size. The figure shows a graph of effect-sizes for analyses of the UK Biobank, Dunedin Study (Dunedin), Brain Genomics Superstruct Project (GSP) and Duke Neurogenetics Study (DNS) samples (solid blue diamonds) and the cross-study effect-size estimated from random-effects meta-analysis (open blue diamond). Gray boxes around the solid-blue diamonds show the weighting of study-specific estimates in the meta-analysis (larger gray boxes indicate higher weights). 95% CIs for estimates are shown as error bars for the study-specific estimates and as the left- and right-extremes of the diamond for the meta-analysis effect-size. The meta-analysis estimate of between-study heterogeneity (I-squared) is listed to the left of the open blue diamond showing the meta-analysis effect-size. The table to the right of the effect-size graph reports values for effect-sizes, 95% CIs, and meta-analysis weights.

**Figure 4.**
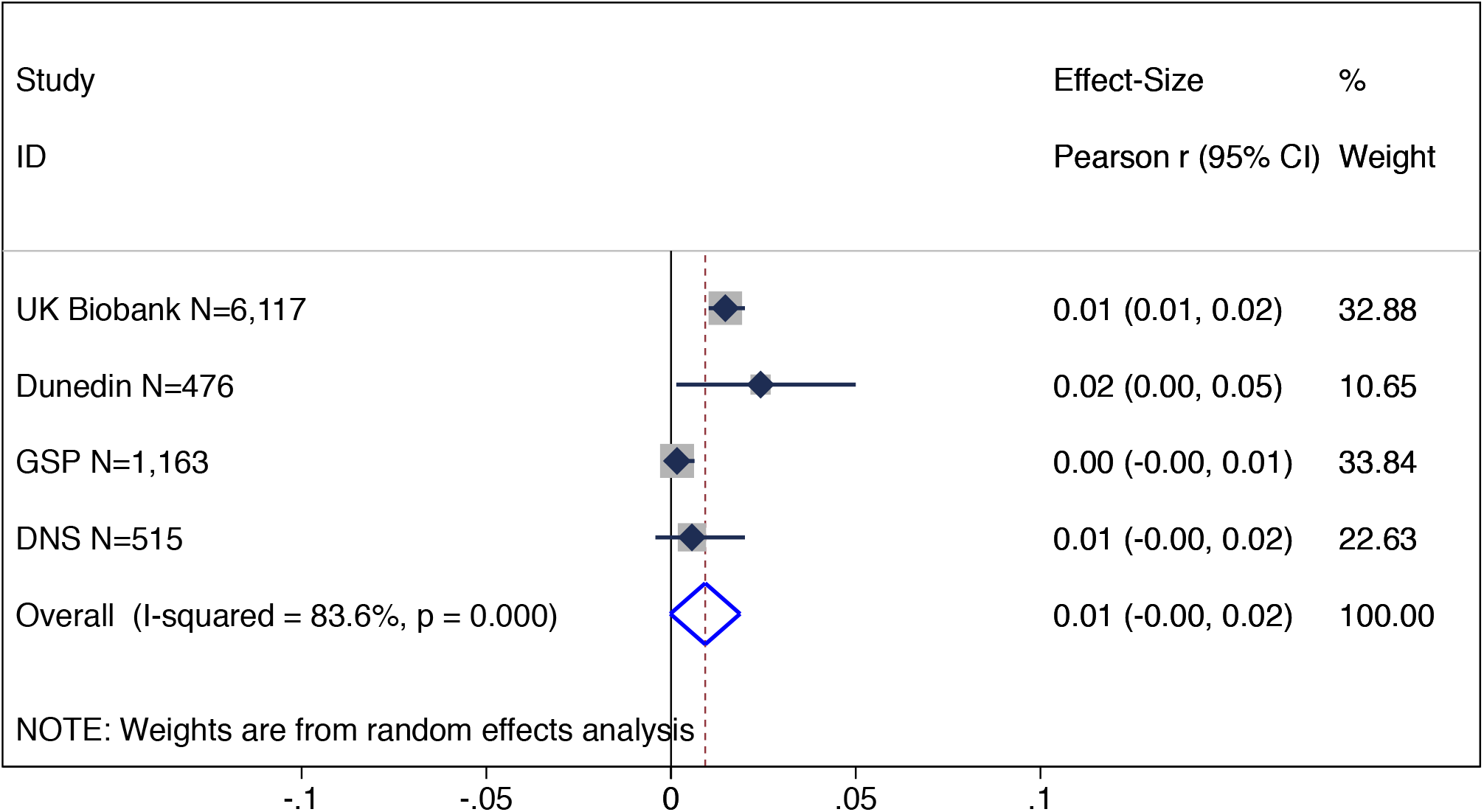
Mediation effect of brain size on the association between the polygenic score for educational attainment and cognitive test scores. The figure shows a graph of effect-sizes for analyses of the UK Biobank, Dunedin Study (Dunedin), Brain Genomics Superstruct Project (GSP) and Duke Neurogenetics Study (DNS) samples (solid blue diamonds) and the cross-study effect-size estimated from random-effects meta-analysis (open blue diamond). Gray boxes around the solid-blue diamonds show the weighting of study-specific estimates in the meta-analysis (larger gray boxes indicate higher weights). 95% CIs for estimates are shown as error bars for the study-specific estimates and as the left- and right-extremes of the diamond for the meta-analysis effect-size. The meta-analysis estimate of between-study heterogeneity (I-squared) is listed to the left of the open blue diamond showing the meta-analysis effect-size. The table to the right of the effect-size graph reports values for effect-sizes, 95% CIs, and meta-analysis weights.

## Acknowledgement

This research was conducted using the UK Biobank Resource (project ID #28174). The Dunedin Multidisciplinary Health and Development Study is supported by the NZ HRC, NZ MBIE, National Institute on Aging grant R01AG032282, R01AG049789, and UK Medical Research Council grant MR/P005918/1. The Duke Neurogenetics Study received support from Duke University as well as US-National Institutes of Health grants R01DA033369 and R01DA031579. MLE is supported by the National Science Foundation Graduate Research Fellowship under Grant No. NSF DGE-1644868. DWB is supported by an early-career fellowship from the Jacobs Foundation. TG is supported by National Institutes of Health grant K99AG054573. AJH is supported by National Institute of Mental Health grant K01MH099232. The authors declare no competing financial interests.

