## Supplementary Materials for "A Polygenic Score for Higher Educational Attainment is Associated with Larger Brains"

**Supplemental**

**United Kingdom Biobank (UK Biobank)**

**Sample.** The UK Biobank is an ongoing study of the determinants of disease in a population-based cohort of volunteers that was recruited from the UK National Health Service records beginning in 2006. The UK biobank is publicly available and described in more detail elsewhere^1,2^. Overall this sample consists of approximately 500,000 UK residents aged 40-69 at recruitment^3^. At the time of writing approximately 10,000 of these subjects have additionally completed MRI imaging^2^. Data used in these analyses were originally downloaded in October 2017 under project ID #28174. In this analysis only self-reported white, non-Hispanic subjects with acceptable MRI segmentation and compliance on the cognitive testing were examined (n = 6117).

**Polygenic Scoring.** Genome-wide genetic data in the UK Biobank was collected on the UK Biobank Axiom array. This array consists of 825,927 markers and was designed to cover common genome-wide genetic variation. Details of the genetic data collection and quality control procedures are described in detail here^4^.

We computed polygenic scores for UK Biobank participants based on the published GWAS of educational attainment^5^. Polygenic scoring was conducted following the method described by Dudbridge^6^ using the PRSice software^7^ package in R version 3.4.0. Briefly, single nucleotide polymorphisms (SNPs) reported in genome-wide association study (GWAS) results were matched with SNPs in the UK Biobank database. For each SNP, the count of phenotype-associated alleles (i.e. alleles associated with more educational attainment) was weighted according to the effect estimated in the GWAS. Weighted counts were summed across SNPs to compute polygenic scores. We used all matched SNPs to compute polygenic scores irrespective of nominal significance for their association with educational attainment.

**Cognitive test performance.** Cognitive ability was measured using 13 verbal-numeric reasoning puzzles completed during a 2-minute time test^8^. Responses were scored as number of correct responses with a maximum score of 13. In the UK biobank data showcase this variable is listed as “fluid intelligence” (<http://biobank.ctsu.ox.ac.uk/crystal/field.cgi?id=20016>) and despite its short administration length has been found to have a moderate Cronbach’s alpha of 0.62^9^. On average subjects used in the analysis answered correctly on just over half of the questions (M = 6.95, SD = 2.10).

**Total Brain Volume.** Image collection details for the UK biobank have been extensively described elsewhere^2^. Briefly, magnetic resonance imaging data were collected on 3 identical 3T Siemens Skyra scanners (General Electric Healthcare, Little Chalfont, United Kingdom) using a 32-channel Siemens head coil. T1-weighted images were obtained using a 3D MPRAGE with the following parameters: TR = 2000 ms; TI = 880 ms; 208 saggital slices, matrix =256×256; slice thickness = 1 mm with no gap; and total scan time = 4 min and 52 s. T1-weighted images were then processed with Sienax^10^ to obtain an estimate of total brain volume (M = 1170.86, SD = 111.14).

**Dunedin.**

**Sample.** The Dunedin Study is a longitudinal investigation of health and behavior in a complete birth cohort.  Study members (N=1,037; 91% of eligible births; 52% male) were all individuals born between April 1972 and March 1973 in Dunedin, New Zealand (NZ), who were eligible based on residence in the province and who participated in the first assessment at age 3.  The cohort represents the full range of socioeconomic status on NZ’s South Island and matches the NZ National Health and Nutrition Survey on key health indicators (e.g., BMI, smoking, GP visits)^11^. The cohort is primarily white; fewer than 7% self-identify as having non-Caucasian ancestry, matching the South Island^11^. Assessments were carried out at birth and ages 3, 5, 7, 9, 11, 13, 15, 18, 21, 26, 32, and 38 years, when 95% of the 1,007 study members still alive took part. Data collection is now ongoing in a brain imaging study of participants, who are now aged 45 years. Currently, brain imaging data are available for 476 study members of white ancestry. This subset of the sample did not differ from the full cohort on mean IQ (mean in subsample = 99, mean in whole sample = 100).

**Polygenic Score.** The Dunedin Study used Illumina HumanOmni Express 12v1.1 BeadChip arrays (Illumina CA, USA) to assay common Single Nucleotide Polymorphism (SNP) variation in the genomes of our cohort members. Additional SNPs were imputed using the impute2 software (version 2.3.1, <https://mathgen.stats.ox.ac.uk/impute/impute_v2.html>;^12^) and 1000 Genomes version-3 reference panel^13^. Imputation was conducted on autosomal SNPs appearing in dbSNP (v140) that were called in >98% of the Dunedin Study samples. Invariant SNPs were excluded. Pre-phasing and imputation were conducted using a 50M base-pair sliding window. The resulting genotype database included genotyped SNPs and SNPs imputed with 90% probability of a specific genotype among the non-Maori members of the Dunedin cohort (n=918). We calculated polygenic scores for the European-descent Dunedin cohort using the PRSice software (v1.22, <http://prsice.info/> ^7^ following the same procedure used in UK Biobank, i.e. we used all matched SNPs to compute polygenic scores irrespective of nominal significance for their association with educational attainment.

**Cognitive Test Performance.** Cognitive testing was conducted with the Wechsler Adult Intelligence Scale-IV (WAIS-IV)^14^ when Study members were aged 38 years. The WAIS-IV is a highly reliable instrument for measurement of general intelligence comprising several tests of different crystalized and fluid cognitive functions. We analyzed general cognitive function measured as full-scale IQ score, which has a population reference mean score of 100 and a standard deviation of 15. Dunedin Study participants in our analysis roughly matched this norm (M=99.49, SD = 15.43).

**Total Brain Volume.** Study participants were scanned using a Siemens Skyra 3T scanner (Siemens Healthcare, Erlangen, Germany) equipped with a 64-channel head/neck coil at the Pacific Radiology imaging center in Dunedin, New Zealand. High resolution structural images were obtained using a T1-weighted MP-RAGE sequence with the following parameters: TR = 2400 ms; TE = 1.98 ms; 208 sagittal slices; flip angle, 9°; FOV, 224 mm; matrix =256×256; slice thickness = 0.9 mm with no gap (voxel size 0.9×0.875×0.875 mm); and total scan time = 6 min and 52 s. T1-weighted images were processed through Freesurfer’s^15^ recon-all imaging pipeline (version 6.0.0) along with T2-weighted flair images to improve the modeling of the pial surface. The BrainSegNotVent value was used as a measure of total brain volume in each subject, which represents the volume of all gray matter and white matter structures in the cortex and cerebellum, excluding the ventricles and brain stem (M = 1218.32 cm^3^, SD = 121.36 cm^3^).

**Brain Genomics Superstruct Project (GSP)**

**Sample.** The GSP sample was collected between 2008 and 2012 in Boston, Massachusetts. Described extensively elsewhere^16^, the sample consists of young adults (18 to 35) primarily recruited through Boston area colleges and universities and Massachusetts General Hospital. Participants came were recruited to be a part of the GSP if they were participating in a study of normal (non-clinical) brain function or serving as a control participant in a case-control study of a clinical population at Massachusetts General Hospital. In this analysis only self-reported white, non-Hispanic subjects with acceptable MRI segmentation (determined by GSP) and compliance on the Shipley IQ were examined (n = 1163).

**Polygenic Scoring.** GSP used two Oragene saliva kits (Oragene, DNA Genotek)

to assay common Single Nucleotide Polymorphism (SNP) variation in the genomes of participants from saliva DNA ^16^. We analyzed genotyped SNPs with MAF > .01, HWE > 1e-6 and missingness < 0.02 using plink v1.90. We computed polygenic scores for European-descent GSP participants based on the published GWAS of educational attainment^5^. Polygenic scoring was conducted following the method described by Dudbridge^6^ using the PRSice software^7^ package in R version 3.4.0 applied in the same way as the UK Biobank sample (described above).

**Cognitive Test Performance.** GSP conducted cognitive testing using the Shipley Institute of Living Scale^17^ to measure full-scale IQ. The Shipley is a quick and robust measure of crystalized ability and fluid reasoning that can be administered online. This measure was scored against a population norm with M=100 and SD=15. GSP subjects showed elevated scores on average with reduced variation relative to the population norms (M = 113.12, SD = 9.02).

**Total Brain Volume.** Imaging details for the GSP have been described elsewhere^16^. Briefly, magnetic resonance imaging data were collected on 3T Tim Trio scanners (Siemens Healthcare, Erlangen, Germany) using a 12-channel head coil. T1-weighted images were obtained using a multi-echo MPRAGE with the following parameters: TR = 2200 ms; TI = 1100ms; TE = 1.5/3.2/5.2/7.0, 144 sagital slices, 1.2 x 1.2 x 1.2 mm; and total scan time = 2 min and 12 s. T1 anatomical images were run with Freesurfer’s recon-all pipeline version 5.3 and T2-weighted images were used to aid segmentation. The BrainSegNotVent value was used as a measure of total brain volume in each subject, which represents the volume of all gray matter and white matter structures in the cortex and cerebellum, excluding the ventricles and brain stem (M = 1174.58, SD = 110.64).

**Duke Neurogenetics Study (DNS)**

**Sample.** The DNS is a large imaging genetics sample (n = 1333) originally designed to investigate the neurogenetic pathways of variation in human behavior among 18- to 22-year-old college students. All participants provided informed consent in accordance with the Duke University Medical Center Institutional Review Board guidelines before participation. All participants were in good general health and free of the following study conditions: (1) medical diagnoses of cancer, stroke, head injury with loss of consciousness, untreated migraine headaches, diabetes requiring insulin treatment, chronic kidney or liver disease; (2) use of psychotropic, glucocorticoid or hypolipidemic medication; and (3) conditions affecting cerebral blood flow and metabolism (e.g., hypertension). The final non-Hispanic Caucasian sample with high-quality MRI, cognitive testing and genetic data consisted of 515 subjects.

**Polygenic Scoring.** DNA was isolated from saliva derived from Oragene DNA self-collection kits (DNA Genotek) customized for 23andMe (www.23andme.com). DNA extraction and genotyping were performed through 23andMe by the National Genetics Institute (NGI), a CLIA-certified clinical laboratory and subsidiary of Laboratory Corporation of America. One of two different Illumina arrays with custom content was used to provide genome-wide SNP data, the HumanOmniExpress or HumanOmniExpress-24^18–21^.

We computed polygenic scores for non-Hispanic Caucasian DNS participants based on the published GWAS of educational attainment^5^. Polygenic scoring was conducted following the method described by Dudbridge^6^ using the PRSice software^7^. Briefly, SNPs reported in GWAS results were matched with SNPs in the DNS database. For each SNP, the count of phenotype-associated alleles (i.e. alleles associated with more educational attainment) was weighted according to the effect estimated in the GWAS. Weighted counts were summed across SNPs to compute polygenic scores. We used all matched SNPs to compute polygenic scores irrespective of nominal significance for their association with educational attainment.

**Cognitive Test Performance.** DNS conducted cognitive testing using the Wechsler Abbreviated Scale of Intelligence^22^. The vocabulary and and matrix reasoning subtests were combined to derive a full-scale measure of IQ, which has a population reference mean score of 100 and a standard deviation of 15. DNS subjects had elevated scores on this scale and reduced variation relative to the population norms (M = 123.73, SD = 7.35).

**Total Brain Volume.** Magnetic resonance imaging data was collected at the Duke-UNC Brain Imaging and Analysis Center using two identical research-dedicated GE MR750 3T scanners (General Electric Healthcare, Little Chalfont, United Kingdom) equipped with high-power high-duty cycle 50-mT/m gradients at 200 T/m/s slew rate, and an eight-channel head coil for parallel imaging at high bandwidth up to 1 MHz. T1-weighted images were obtained using a 3D Ax FSPGR BRAVO with the following parameters: TR = 8.148 ms; TE = 3.22 ms; 162 axial slices; flip angle, 12°; FOV, 240 mm; matrix =256×256; slice thickness = 1 mm with no gap; and total scan time = 4 min and 13 s.

T1-weighted images were then processed through Freesurfer’s^15^ recon-all (version 6.0.0) imaging pipeline. Specifically, the BrainSegNotVent value was used as a measure of total brain volume in each subject, which represents the volume of all gray matter and white matter structures in the cortex and cerebellum, excluding the ventricles and brain stem (M = 1162.40, SD = 110.34).

**Statistical Analyses.** We analyzed associations using linear regression models. All models were adjusted for sex. Models including the polygenic score were adjusted for the first 10 principal components estimated from the genome-wide SNP data to account for residual population stratification within the European-descent samples analyzed^23^. Models of UK biobank and GSP data were adjusted for age. (The Dunedin Study is a single-year birth cohort and DNS participants vary in age only by a year or so). Analysis of individual studies were conducted in R (version 3.4.0). Linear regressions were performed using the lm function. Mediation analyses were performed using a system of equations approach^24^ implemented with the *mediation* package^25^ in R, using nonparametric bootstrapping with 1000 iterations. We combined estimates across studies using random effects meta-analysis^26^ implemented using the STATA software (version 15).

**Supplemental Tables**

**Supplemental Table S1.** Effect-size estimates (standardized regression coefficients interpretable as Pearson r) for analysis in the UK Biobank, Dunedin Study, Brain Genomics Superstruct Project(GSP, and Duke Neurogenetics Study (DNS) samples.

|  | r | SE | P-value |
| --- | --- | --- | --- |
| Polygenic Score associations with Cognitive Performance | | | |
| UK Biobank | .20 | .01 | <.001 |
| Dunedin | .28 | .05 | <.001 |
| GSP | .19 | .03 | <.001 |
| DNS | .05 | .04 | .220 |
| Total Brain Volume associations with Cognitive Performance | | | |
| UK Biobank | .21 | .02 | <.001 |
| Dunedin | .35 | .06 | <.001 |
| GSP | .12 | .04 | .002 |
| DNS | .16 | .06 | .004 |
| Polygenic Score associations with Total Brain Volume | | | |
| UK Biobank | .08 | .01 | <.001 |
| Dunedin | .08 | .04 | .033 |
| GSP | .02 | .02 | .380 |
| DNS | .04 | .03 | .288 |

**Supplemental Table S2.** Effect sizes and confidence intervals for total, direct, and indirect (mediated) effects estimated in mediation analysis of UK Biobank, Dunedin Study, Brain Genomics Superstruct Project (GSP), and Duke Neurogenetics Study (DNS) samples. The indirect effect estimates test the hypothesis that total brain volume mediated the association between educational attainment polygenic score and cognitive test performance. Effect-sizes are shown with 95% Confidence intervals estimated from 1000 bootstrap replications in brackets.

|  | Total Effect | Direct Effect | Indirect Effect |
| --- | --- | --- | --- |
| UK Biobank | .20 [.17, .22] | .18 [.16, .21] | .01 [.01, .02] |
| Dunedin Study | .28 [.21, .36] | .26 [.19, .33] | .02 [.00, .05] |
| GSP | .18 [.12, .23] | .18 [.12, .23] | .00 [.00, .00] |
| DNS | .05 [-.04, .14] | .05 [-.04, .14] | .01 [.00, .02] |

**Supplemental Table S3.** Sensitivity analysis results. Table shows effect-sizes from analysis of the entire UK Biobank sample (N=5701) and the UK Biobank sample restricted to participants with high cognitive test scores (N=1401).

|  | UK Biobank  r (SE) | UK Biobank High Cognitive Performance r (SE) |
| --- | --- | --- |
| Cognitive Performance M(SD) | 7.0 (2.1) | 9.8 (0.9) |
| Polygenic Score-> Cognitive Performance | .20 (.01) | .09 (.03) |
| Total Brain Volume -> Cognitive Performance | .21 (.02) | .12 (.03) |
| Polygenic Score -> Total Brain Volume | .08 (.01) | .05 (.02) |

**Supplemental Table S4.** Characteristics of UK Biobank participants in the high cognitive performance sample analyzed in sensitivity analysis.

| Sample | Sample Size | Age (SD) | Sex(%female) | Cognitive test score (SD) | TBV (cm^3^) (SD) |
| --- | --- | --- | --- | --- | --- |
| High Cognitive Performance UK Biobank | 1401 | 60.75(6.75) | 44.07 | 9.8 (.92) | 1203.34(105.08) |
